# *APOE2* promotes longevity independent of Alzheimer’s disease

**DOI:** 10.1101/2020.08.31.255141

**Authors:** Mitsuru Shinohara, Takahisa Kanekiyo, Masaya Tachibana, Aishe Kurti, Motoko Shinohara, Yuan Fu, Jing Zhao, Xianlin Han, Patrick M. Sullivan, William G Rebeck, John D. Fryer, Michael G. Heckman, Guojun Bu

**Author notes:** Correspondence: Mitsuru Shinohara, Ph.D., Department of Aging Neurobiology, National Center for Geriatrics and Gerontology, 7-430 Morioka, Obu, Aichi, 474-8511, Japan. & Guojun Bu, Ph.D., Department of Neuroscience, Mayo Clinic, 4500 San Pablo Road, Jacksonville, FL 32224, USA.

## Abstract

**Objective:** Although apolipoprotein E (*APOE*) allele associates with longevity, its mechanism is not understood. The protective effects of *APOE*2 and the deleterious effects of *APOE4* on Alzheimer’s disease (AD) risk may confound *APOE* effects on longevity.

**Methods:** We analyzed a large number of subjects from the National Alzheimer’s Coordinating Center (NACC), and animal models expressing human apoE isoforms in the absence of AD.

**Results:** Clinically, the *APOE2* allele was associated with longer lifespan, while *APOE4* associated with shorter lifespan, compared to the common *APOE3* allele. This effect was also seen irrespective of clinical AD status, and in subjects with little amyloid pathology or after adjustment for AD-related pathologies. In animal studies, apoE2-TR mice also exhibited longer lifespan, while apoE4 showed some trends of shorter lifespan. Notably, old apoE2-TR mice kept activity measured by open field assay, associated with longer lifespan. Evidence of preserved activity in *APOE2* carrier was also obtained in clinical records. In animal studies, higher levels of apoE2 in brain and plasma were correlated with activity. Moreover, lower levels of total cholesterol in the brain and higher levels of high-density lipoprotein cholesterol and triglycerides in the plasma of apoE2-TR mice were associated with apoE levels and more activity.

**Interpretation:** *APOE2* can contribute to longevity independent of AD. Preserved activity would be an early-observable feature of apoE2-mediated longevity, where higher levels of apoE2 and its-associated lipid metabolism might be involved.

## Introduction

It is well established that the ε4 allele of the apolipoprotein E gene (*APOE4*) is a strong risk factor for late-onset Alzheimer’s disease (AD) and that *APOE2* is protective (*1*). Additionally, both *APOE* alleles are also associated with longevity. Several case-control studies have shown higher frequencies of *APOE2* in elderly individuals and centenarians compared to younger populations, whereas the frequency of *APOE4* is lower in the older individuals (*2-4*). Some longitudinal studies have also demonstrated beneficial effects of *APOE2* and deleterious effects of *APOE4* on longevity (*5, 6*). Effects of *APOE* on longevity have also been observed in unbiased GWAS analyses (*7, 8*). Despite the accumulated evidence regarding *APOE* effects on longevity, the underlying mechanism has rarely been addressed.

ApoE proteins form lipoprotein particles and regulates lipid transport in both the central nervous systems and periphery. While apoE isoforms can directly affect Alzheimer’s neuropathology, including the accumulation of amyloid-β (Aβ) and tau, increasing evidence has demonstrated that apoE isoforms also contribute to cognitive function through AD neuropathology-independent pathways (*1, 9*). These effects might be mediated by apoE-regulated lipid metabolism, synaptic function, vascular integrity, and/or neuroinflammation. Recently, by performing both clinical and preclinical analysis, we showed that *APOE2* protects against age-associated cognitive decline independent of AD neuropathology. Interestingly, these effects were also independent of synaptic and neuroinflammatory changes but associated with cholesterol metabolism in both the brain and periphery (*10*).

In this study, we aimed to address the effects of *APOE* on longevity and assess their relationship with AD, including clinical symptoms and neuropathology. Toward this, we analyzed clinical records of a large number of subjects enrolled by the National Alzheimer’s Coordinating Center (NACC). In addition, we analyzed mouse models expressing each human apoE isoform under the control of the mouse endogenous promoter (targeted replacement or “TR” mice). Our results provide important insights into how *APOE2* contributes to longevity.

## Materials and Methods

### Human clinical and neuropathological data

The clinical data from NACC, which were collected by the 34 past and present Alzheimer’s Disease Centers (ADCs) from September 2005 to November 2016 as the longitudinal Uniform Data Set (*11*), were assessed in this study. We restricted our analysis to 24,661 subjects whose sex, race, and *APOE* genotypes were available in order to assess the effects of *APOE* on longevity (∼70% of total subjects). NACC subjects are regarded as a referral-based or volunteer series, and the majority of these subjects are Caucasians (84.2%) with variable cognitive status at final visit (36.0% normal cognition, 4.2% impaired but not MCI, 15.6% MCI, 44.3% dementia) (Supplementary Table 1). This cumulative database also contains neuropathological data from a subset of deceased individuals (N=3,528, 65.2% of total deceased subjects). For comparison with the *APOE* ε3/ε3 genotype (i.e. *APOE3*) in the primary analysis, *APOE* ε2/ε2 and ε2/ε3 genotypes were grouped together into an *APOE2* group, while *APOE* ε3/ε4 and ε4/ε4 genotypes were grouped together into an *APOE4* group. Although not the primary focus of this study, comparisons between *APOE* ε2/ε4 and ε3/ε3 genotypes were also made, as shown in supplementary tables.

NACC demographic variables that were utilized in our study included death, age at death, age at initial visit, age at final visit, sex, and race. Degree of cognitive impairment was assessed by cognitive status at final visit using the NACCUDSD variable and also by clinically diagnosed AD at final visit. Clinically diagnosed AD at final visit was defined as presence of both dementia on NACCUDSD, and a value of “yes” on NACCALZD. Cardiovascular factors (occurrence of these at any visit was assessed) that were measured included hypertension, transient ischemic attack, pacemaker, angioplasty/endarterectomy/stent, heart attack/cardiac arrest, cardiac bypass procedure, atrial fibrillation, hypercholesterolemia, congestive heart failure, and stroke. We also utilized data from the geriatric depression scale (GDS), where answers to 15 different yes/no questions are recorded as well as a total GDS score that ranges from 0 to 15 and represents the sum of the scores for the individual GDS questions.

Neuropathological information was collected regarding Consortium to Establish a Registry for Alzheimer’s Disease (CERAD) diffuse plaque score, CERAD neuritic plaque score, Braak NFT stage, and the presence of vascular pathology. It is of note that the neuropathologically-assessed cohort had a different frequency of dementia and AD, compared to the overall cohort, with much higher rates of dementia (78.3% vs. 44.3%) and AD (58.0% vs. 35.3%) at final NACC visit, likely due to higher rate of postmortem analysis of demented subjects (Supplementary Table 2). Minimal amyloid pathology was defined as both no neuritic plaques and no or sparse diffuse plaques (*10*).

### Statistical analysis of NACC human clinical data

Statistical analyses were performed using SAS (version 9.4; SAS Institute, Inc., Cary, North Carolina) and R Statistical Software (version 3.2.3; R Foundation for Statistical Computing, Vienna, Austria). The Kaplan-Meier method was used to estimate survival (i.e. after birth), where censoring occurred at the date of the last NACC visit for subjects who did not die. In the primary analysis including all subjects, Cox proportional hazards regression models were used to compare survival between *APOE* carriers. Models were first adjusted for sex and race, and then were subsequently adjusted for cognitive status at final NACC visit and clinically diagnosed AD at final NACC visit in order to account for these two strong confounding factors that are related to both *APOE* genotype and survival. We also stratified by the AD/non-AD status and compared survival using Cox regression models that were adjusted for sex and race. In a sensitivity analysis, we additionally adjusted our Cox models for the aforementioned ten cardiovascular factors in the subset (97.7%) of subjects who had this information available.

We subsequently examined only neuropathologically assessed cases in order to evaluate the relationship between *APOE*-mediated longevity and AD-related neuropathology. Specifically, Cox proportional hazards regression models that were adjusted for sex, race, CERAD diffuse plaque score, CERAD neuritic plaque score, Braak NFT stage, and the presence of vascular pathology were utilized to compare survival between *APOE* carriers. These comparisons were also made separately in subjects with minimal amyloid pathology, where Cox proportional hazards regression models were adjusted for sex and race. Finally, additional adjustment for cardiovascular factors in Cox regression models was examined in a sensitivity analysis.

In order to adjust for the fact that two primary comparisons of survival were made (i.e. between *APOE3* and *APOE4* subjects, and also between *APOE3* and *APOE2* subjects), we applied a Bonferroni correction after multiple testing, after which p-values <0.025 were considered as statistically significant. All statistical tests were two-sided.

For analysis of the GDS individual items and total score, we restricted the analysis to subjects who were cognitively normal at their last evaluation in order to exclude the possibility of obtaining false answers from cognitively impaired subjects. Comparisons between *APOE3* and *APOE4* subjects, and also between *APOE3* and *APOE2* subjects, were made using binary logistic regression models for the 15 individual yes/no items, and using proportional odds logistic regression models for the ordinal 0-15 total GDS score (where total GDS scores ≥10 were collapsed together into one category due to their low frequencies). Models were adjusted for sex, race, and age at the time of the GDS questionnaire. Odds ratios (ORs) and 95% CIs were estimated and are interpreted as the multiplicative increase in the odds of presence of the given characteristic for analysis of individual GDS items, and as the multiplicative increase in the odds of a higher score for analysis of the total GDS score. The primary comparison was regarding the GDS item “have you dropped many of your activities and interests”. In secondary analysis, comparisons of the remaining GDS items as well as the GDS score were also made to examine whether *APOE* genotype is systematically associated with depression rather than with the more specific “dropped activities and interests” GDS question. In order to adjust for the two comparisons of each GDS item, we applied a Bonferroni correction separately for each GDS outcome measure, after which p-values <0.025 were considered as statistically significant. Additionally, we evaluated the association between the GDS “dropped activities and interests” item and survival using a Cox proportional hazards regression model that was adjusted for sex, race, and *APOE* genotype group, where the same group of subjects who were cognitively normal and older than 60 at the time of GDS evaluation were examined. Again, censoring occurred at the date of the last NACC visit for subjects who did not die. All statistical tests were two-sided.

### Animals

Human apoE TR mice expressing human apoE2, apoE3, or apoE4, under the control of the mouse apoE promoter, known as apoE-targeted replacement (apoE-TR) (*12-14*), and *Apoe*-knockout (KO) mice (*15*) on a pure C57BL/6 background were used in this study. All cohorts examined in this study were generated from homozygous breeding pairs, group housed without enrichment structures in a specific pathogen-free environment in ventilated cages and used in experiments according to the standards established by the Mayo Clinic Institutional Animal Care and Use Committee (IACUC). We analyzed two animal cohorts in this study (Supplementary Table 4). Sample size was determined according to the previous publications related to mouse longevity study (*16*) as well as the capacity of our animal facility. The first cohort was employed for the survival analysis, and the second was employed for biochemical analyses. In the first cohort used for the survival analysis (n=118), we euthanized mice when they showed severe age-related symptoms (33.9% of total animals) including severe weakness (a severely hunched posture, rapid weight loss (10% decrease in one week), and lethargy), large abdominal masses, significant skin lesions, and prolapse, according to the guidelines of the Mayo Clinic IACUC to avoid severe pain and distress before death in animals. These symptoms were diagnosed by the expert on-site veterinarian, according to the NIH Body Condition Scoring system (*17*). Two mice (2% of the total mice) were also euthanized for reasons not related to these health issues; the animals survived until the end of our project, and thus, they were censored in the whole survival analysis. The remaining 66.1% of mice died naturally. A significant difference in the number of euthanized/naturally-dead mice was not observed among *APOE* genotype groups. In the second cohort employed for biochemical analyses, we analyzed the same apoE2-, apoE3-and apoE4-TR mice as we have previously reported (*10*), except *Apoe*-KO mice were also included. All experiments were conducted in a blinded manner, and all date were included in the analyses without defining outliers nor exclusion criteria.

### Behavioral testing

All tests were performed during the first half of the light cycle. All mice were acclimated to the testing room for 1–2 hours before testing. All behavioral equipment was cleaned with 30% ethanol between each animal. All mice were returned to their home cages and home room after each test.

### Open-field assay

Mice were placed in the center of an open-field area (40 × 40 x 30 cm, width x length x height) and allowed to roam freely for 15 minutes. Side-mounted photobeams raised 7.6 cm above the floor were used to measure rearing while an overhead camera was used to track movement including total distance traveled, average speed, and time spent mobile, with AnyMaze software (Stoelting Co, Wood Dale, IL, USA).

### Elevated plus maze test

As a formal test of anxiety/exploration, the entire maze is elevated 50 cm above the floor and consists of four arms (50 × 10 cm) with two of the arms enclosed with roofless gray walls (35 × 15 cm, L × H). Mice were tested by placing them in the center of the maze facing an open arm, and their behavior was tracked for 10min using an overhead camera and AnyMaze software.

### Rotarod

Motor coordination was measured using an automated rotarod system (Rotamex-5 Columbus Instruments). The spindle dimensions were 3.0cm × 9.5cm and the speed of the rod was set to 4–40rpm acceleration, increasing 1rpm every 5 seconds. The apparatus was equipped with a sensor that automatically stops the timer if the mice fall off the rod. Mice were trained for 4 days in 2 consecutive trials per day, allowing 10 minutes of rest between trials.

### Tissue harvest and sample preparation

We harvested mice and prepared samples for biochemical analyses as previously described (*10*).

### ELISAs and other biochemical assays

Levels of apoE, postsynaptic density 95 (PSD95), glial fibrillary acidic protein (GFAP), CD11b, tumor necrosis factor α (TNFα) and interleukin-1β (IL1β), and cholesterol in the brain were determined, as previously described (*10*). Levels of C-reactive protein (CRP) were determined using a commercial ELISA kit (R&D Systems). Plasma levels of total cholesterol, HDL-cholesterol and triglycerides were determined using enzymatic assays, according to the manufacturer’s instructions (Wako). Plasma levels of non-HDL-cholesterol were determined by subtracting HDL-cholesterol levels from total cholesterol levels in the plasma (*18*).

### Statistical analysis of animal data

All statistical analyses were performed using JMP Pro software (version 12, SAS Institute Inc.). Log-rank tests that were adjusted for sex were used to compare survival between *APOE* genotype groups; p-values <0.0083 were considered as statistically significant after applying a Bonferroni correction for multiple comparisons for the six pair-wise comparisons that were performed. Given evidence of a different survival pattern for apoE3-TR mice over time in comparison to the other groups, specifically with dramatically different pattern after day ∼800, we also made pair-wise comparisons of “early survival” with the apoE3-TR group by censoring all survival times at day 800; p-values <0.0167 were considered as statistically significant after applying a Bonferroni correction for multiple comparisons for the 3 additional comparisons that were performed.

For behavior experiments and biochemical experiments, comparisons of outcomes across *APOE* genotypes were made (1) by linear regression models adjusting for sex followed by Tukey’s HSD test when both genders were included, and (2) one-way ANOVA followed by Tukey-Kramer test when only one gender was included. Comparisons between young and old mice within each *APOE* genotype group were made by linear regression models adjusting for sex followed by pair-wise two-sided Student’s t tests. To assess the association between age at death, behavior experiment results, and/or biochemical measurements, the Pearson correlation test was conducted to calculate correlation coefficients (r) and p-values.

Details of statistical tests are also provided in each of the Figure legends. Unless otherwise noted, p-values less than 0.05 were considered significant after appropriate adjustment for multiple comparisons.

## Results

### *APOE* is associated with longevity independent of AD in clinical cohorts

To evaluate relationships of each *APOE* allele with longevity, we analyzed NACC clinical records that collected clinical and neuropathology data of large number of demented and non-demented people in a longitudinal manner (*11*). In NACC subjects whose *APOE* genotype, sex, and race were available (n=24,661, demographics shown in Supplementary Table 1), 5,413 (21.9%) of subjects were recorded as dead. We analyzed their survival after birth. As shown in Fig. 1A, compared to subjects with ε3/ε3 genotype (i.e. “*APOE3*”), those who were ε3/ε4 or ε4/ε4 (i.e. “*APOE4*”) had worse survival, while individuals who had ε2/ε2 or ε2/ε3 genotype (i.e. “*APOE2*”) had better survival. Notably, these effects were persisted in analyses that adjusted for sex, race, AD, cognitive status, and cardiovascular factors in all subjects (“All subjects” in Fig. 1B). Moreover, stratifying non-AD and AD status keep these *APOE* effects significant (“No AD” and “AD” in Fig. 1B).

**Fig. 1.**
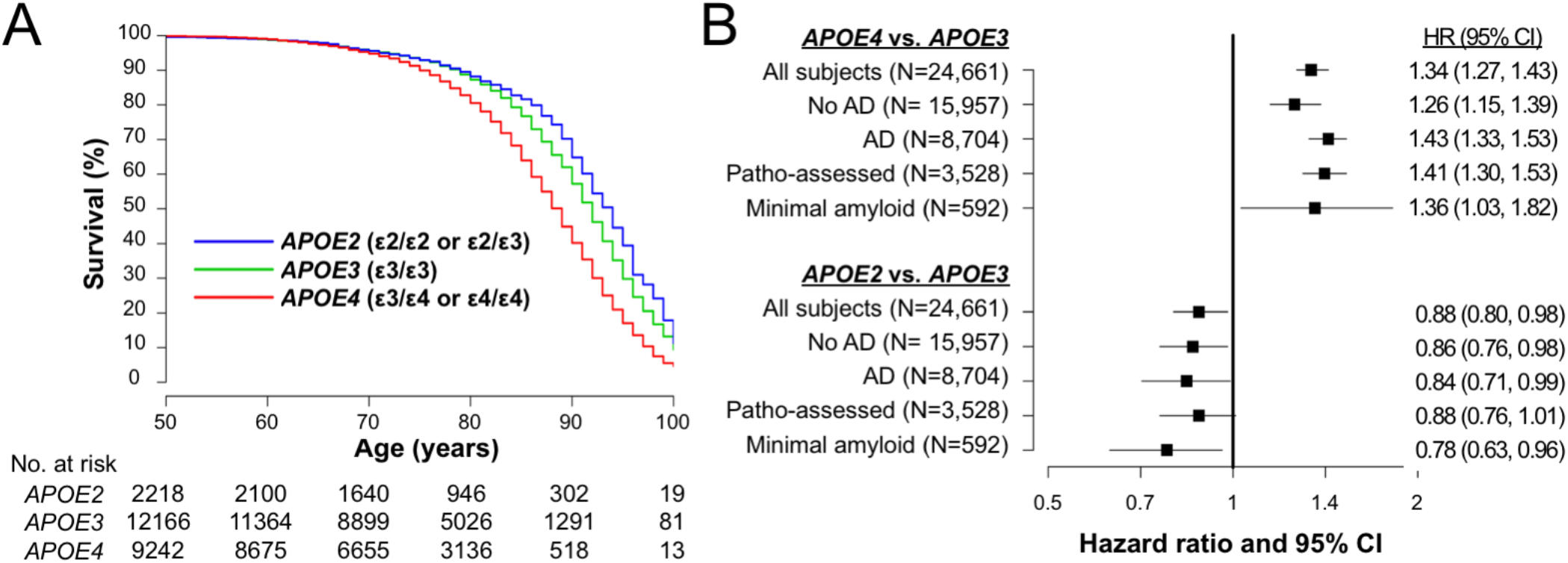
*APOE* associate with lifespan irrespective of AD in clinical cohorts. (A) Kaplan-Meier survival curve according to *APOE* genotype. (B) Effects of *APOE4* (ε3/ε4 or ε4/ε4) or *APOE2* (ε2/ε3 or ε2/ε2) on survival compared to *APOE3* (ε3/ε3) as a reference in all subjects (“All subjects”), when stratifying by AD diagnosis at last visit (“No AD” or “AD”), in subjects with neuropathologically-assessment (“Patho-assessed”), and subjects with minimal amyloid pathology (“Minimal amyloid”). HR=hazard ratio. CI=confidence interval. HRs and 95% CIs result from Cox proportional hazards regression models. Models for all subjects were adjusted for sex, race, cognitive status at last visit, presence of AD at last visit, and cardiovascular factors. The models for subjects with neuropathological assessment were adjusted for sex, race, CERAD diffuse plaque score, CERAD neuritic plaque score, Braak NFT stage, presence of vascular pathology, and cardiovascular factors. All other models were adjusted for sex, race and cardiovascular factors.

We subsequently focused only on the subgroup of 3,528 subjects who were neuropathologically assessed after their death (demographics shown in Supplementary Table 2) in order to evaluate the relationships between *APOE* and longevity while accounting for AD-related neuropathology. When adjusting for sex, race, diffuse plaque score, neuritic plaque score, NFT stage, the vascular pathology, and cardiovascular factors, survival was again worse for *APOE4* subjects and better for *APOE2* subjects compared to the *APOE3* group (“Patho-assessed” in Fig. 1B). Moreover, the harmful effects of *APOE4* and the beneficial effects of *APOE2* on mortality were even observed in individuals without remarkable amyloid pathology (“Minimal amyloid” in Fig. 1B).

These results were precisely described in Supplementary Table 3. Although not of primary interest, ε2/ε4 genotype was mildly (but not significantly) associated with worse survival compared to ε3/ε3 (Supplementary Table 3). These results indicate that *APOE* contributes to longevity independent of clinical and neuropathological status of AD. However, as in any epidemiologic study, unmeasured confounding variables could have affected our results.

### *APOE2* benefits lifespan in animal models, associated with preserved activity

We thus analyzed human apoE-targeted replacement (apoE-TR) (*12-14*) and *Apoe*-knockout (*15*) mice to examine whether the effects of *APOE* on lifespan can indeed be observed in the absence of AD. In the absence of overexpressing mutant *APP* or mutant *MAPT* gene, these mice do not display remarkable amyloid or tau deposition such as those seen in AD (*10, 19, 20*). In the survival mouse cohort (n=118), due to the regulation of our animal experiment, we euthanized mice (33.9% of total animals) when they showed severe age-related symptoms including severe weakness, masses, significant skin lesions etc. that were diagnosed by the expert on-site veterinarian to avoid severe pain and distress before death (see Materials and Methods), while there was no difference in *APOE* effects on these severe symptoms (Supplementary Table 4). The remaining 66.1% of mice died naturally. During their lives, we performed a non-invasive battery of behavioral tests including the open-field assay (OFA) and elevated plus maze (EPM) at both young (4∼7 months age) and old (21∼24 months age) ages, and rotarod test at old age (Fig. 2A). Overall, as shown in Fig. 2B, apoE2-TR mice showed a reduced risk of all-cause mortality when compared to apoE3-TR (P=0.0004), apoE4-TR (P=0.007), and *Apoe*-KO (P=0.0001). Notably, as the survival pattern for “control” apoE3-TR mice varied over time, we also made pair-wise comparisons of “early survival” with the apoE3-TR group by censoring all survival times at day 800. For these comparisons of “early survival”, we observed worse survival for apoE4-TR (P=0.013). Additionally, the median age at death was in the order of apoE2-TR (911 days) > apoE3-TR (825 days) > apoE4-TR (753 days) > *Apoe*-KO (738 days), where apoE4-TR mice showed ∼10% decrease in median lifespan, compared to apoE3-TR mice (Fig. 2B). Nonetheless, these animal model data support beneficial effects of *APOE2* on mortality in the absence of AD.

**Fig. 2.**
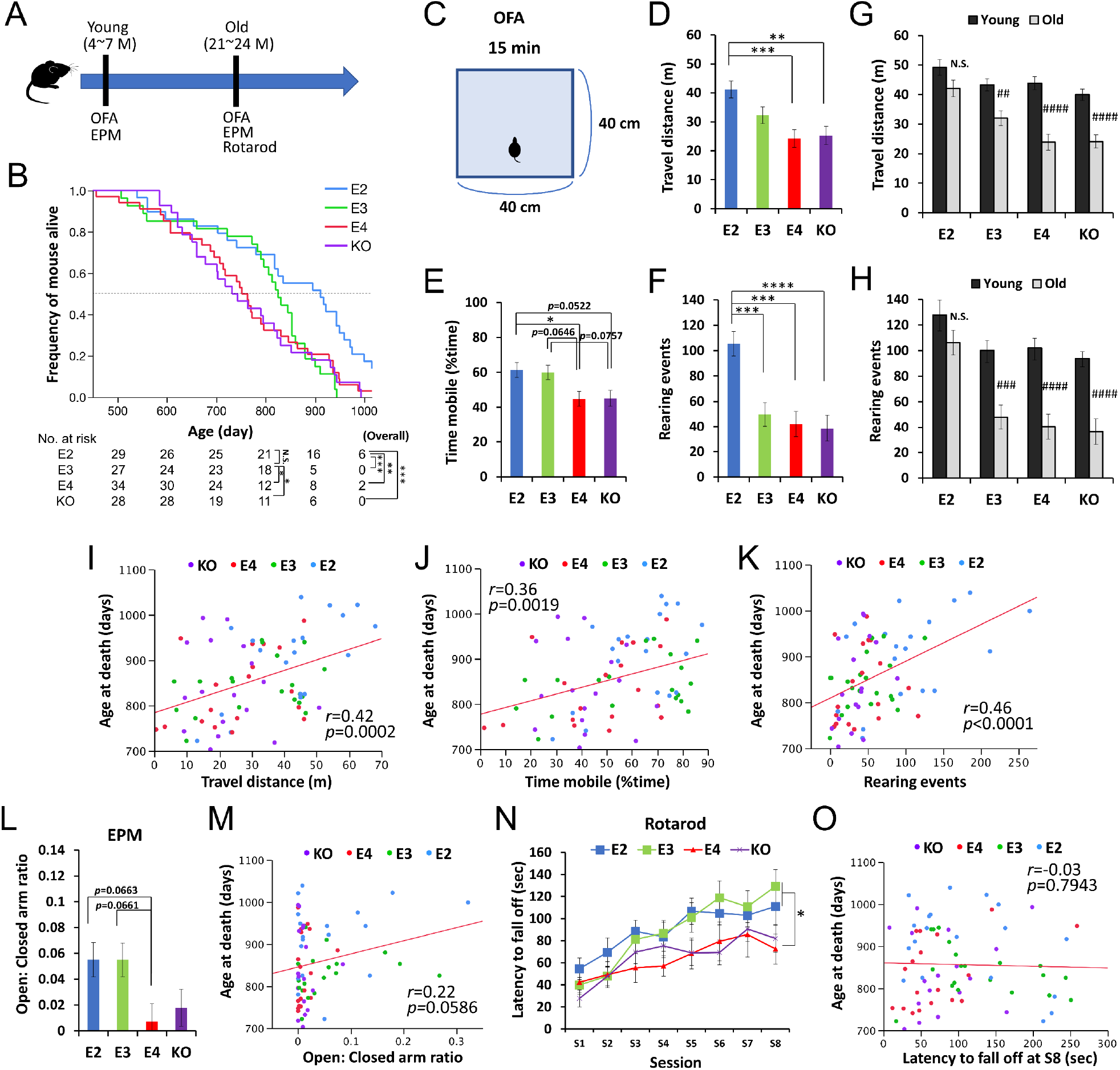
*APOE2* benefits lifespan and preserved activity levels in animal models. (A) The timeline of the survival cohort. The OFA and EPM were performed at both young (4∼7 months) and old (21∼24 months) ages. Rotarod test was also performed at old age. (B) Kaplan-Meier survival curve of apoE-TR and *Apoe*-KO mouse models (n = 118), categorized by *APOE* carrier status with number of mice at risk. *p<0.05, **p<0.01, ***p<0.001 after Bonferroni correction for multiple comparisons; calculated by Cox proportional hazards regression models that were adjusted for sex. Median survival for each group is the point at which the Kaplan-Meier survival curves intersects with the dotted horizontal line. (C) Diagram of OFA. (D-I) Total distance traveled (D & G), percent time spent mobile (E), and number of rearing events (F & H) in the OFA were compared among *APOE* genotype groups at old age (D-F) or between young and old ages within the same *APOE* genotype (G-H) after adjusting for sex (n = 17-30 mice/group). (I-J) Total distance traveled (I), percent time spent mobile (J), and number of rearing events (K) in the OFA performed at old age are plotted against the age at death. (L & M) Ratios of time stayed at open arm versus closed arm measured by EPM test at old age were compared among *APOE* genotype groups after adjusting for sex (L), and plotted against following age at death (M) (n = 17-30 mice/group). (N & O) Latency to fall off at each session of rotarod test at old age was compared among *APOE* genotype groups after adjusting for sex (N), and plotted against following age at death (O) (n = 17-20 mice/group). Data are presented as adjusted means ± standard errors of the means. *p<0.05, **p<0.01, ***p<0.001, ****p<0.0001; compared among *APOE* genotype groups using the Tukey’s HSD test (D-F & L) or repeated-measures one-way ANOVA followed by Tukey-Kramer test (N)., or #p<0.05, ##p<0.01, ###p<0.001, ####p<0.0001; compared with young mice using two-sided Student’s t test (G-H). (I-K, M & O) Correlation coefficients (r) and p-values were calculated using the Pearson correlation test. N.S. = not significant.

In the OFA (Fig. 2C) performed at old age, apoE2-TR mice generally showed greater locomotor and exploratory activity, compared to other mice (Fig. 2D-F). In particular, number of rearing events was significantly increased in apoE2-TR mice even when compared to “control” apoE3-TR mice (Fig. 2F). Notably, the effect of *APOE* genotype on these activity levels was more prominent for the number of rearing events (P<0.0001), which better reflected the exploratory behavior, than the distance traveled (P=0.0005) or the time spent mobile (P=0.0056). Moreover, when comparing the scores at old age with the ones performed at young age, we observed that apoE2-TR mice showed less of an age-associated decrease in these activities (Fig. 2G&H). More interesting, significant correlations were observed between the activity levels in the OFA and age at death across *APOE* genotypes (Fig. 2I-K). There were no significant changes in their body weights of this cohort (data not shown). We also observed some trends of positive association (though not significant, P=0.059) between age at death and results of EPM test (Fig. 2L&M), which primarily measure the exploratory/anxiety phenotype, though no correlations for rotarod test that purely measure locomotive ability (Fig. 2N&O).

### *APOE2* is associated with preserved activity in clinical cohorts

These results prompted us to re-review the NACC records to examine the clinical relevance of *APOE* effects on activity. As a candidate variable that might reflect the results observed in the animal experiments, we searched for “activity” in NACC variables, and identified “dropped activities and interests” as one component of the geriatric depression scale (GDS) test. Despite its subjective nature, previous studies suggest that this question could reflect age-related changes of some form of physical activity independent of depression (*21, 22*). As these GDS components could be confounded by cognitive status, we analyzed subjects who were cognitively normal at their final GDS evaluation. We also assessed only subjects who were over 60 years old in order to assess age-related changes of this measure, resulting in a sample size of 9,530 subjects. Compared to *APOE3* subjects, we observed a lower frequency of “dropped activities in interests” for *APOE2* subjects (Odds ratio [OR]=0.80, P=0.019) in analysis that was adjusted for sex, race, and age, while there was no significant difference between *APOE3* and *APOE4* subjects (OR=0.98, P=0.82) (Fig. 3A, top). Subsequently, comparisons of the remaining 14 GDS items as well as the GDS score were also made between the three *APOE* groups in order to evaluate whether *APOE* genotype is systematically associated with depression. We observed no such evidence; total GDS score was not significantly different in comparison to *APOE3* subjects for either *APOE2* subjects or *APOE4* subjects (Fig. 3A, middle), and “feeling helpless” was the only other individual GDS item that differed significantly between *APOE2* and *APOE3* subjects (OR=0.69, P=0.005) (Supplementary Table 5). Notably, consistent with previous findings (*10, 23, 24*), the risk of “problems with memory” was significantly higher for *APOE4* compared to *APOE3* subjects (OR=1.18, P=0.020) (Fig. 3A, bottom), supporting the validity of our other GDS findings. Moreover, in exploratory analysis using “dropped activities and interest” as a marker of lifetime activity level in this same cohort of 9,530 subjects, we observed that “dropped activities and interests” was associated with a shorter lifespan in analysis adjusted for sex, race, and *APOE* genotype (HR=1.20, P=0.010) (Fig. 3B). These clinical data are consistent with the animal data indicating that some form of activity in the lifetime is associated with *APOE2*-mediated longevity.

**Fig. 3.**
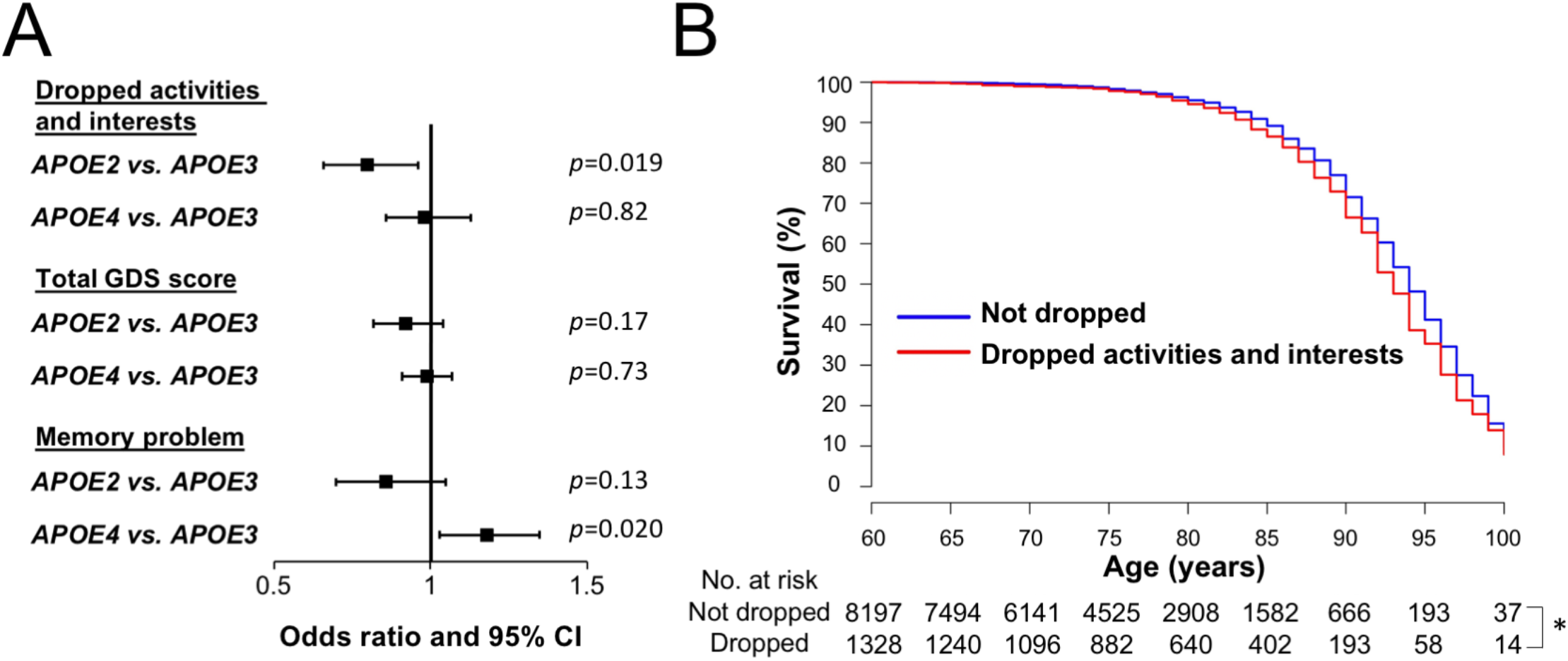
*APOE2* is associated with preserved activity in clinical cohorts. (A) Odds ratio with 95% CIs of *APOE2* or *APOE4* on “dropped activities and interests”, “total GDS score”, and “memory problem” compared to *APOE3*, as calculated by logistic regression models that were adjusted for sex, race, and age at the time of the GDS questionnaire. (B) Kaplan-Meier survival curve of subjects with/without “dropped activities and interests”; dropped activities and interests were associated with poorer survival (Hazard ratio=1.20, P=0.010). *p<0.05; calculated by Cox proportional hazards regression analysis adjusting for sex, race, and APOE genotype group.

### Activity associates with apoE and lipid levels in animal models

We then analyzed biochemical changes in the CNS and periphery of an independent mouse cohort that was harvested within one month of the OFA test (Fig. 4A, young group = 5∼7 months of age, old group = 21∼24 months of age, Supplementary Table 4). Consistent with the survival cohort, old apoE2-TR mice had a greater ability to retain their activity levels in the OFA (Fig. 4B-D). The relative levels of apoE in the brain, CSF and plasma were in the order of apoE2-TR > apoE3-TR > apoE4-TR, as previously reported (*10*). Activity levels in the OFA correlated with such different apoE levels in these tissues of old mice across *APOE* genotypes (Fig. 4E-H & Supplementary Table 6). We analyzed changes in synaptic or inflammatory markers, including PSD95, GFAP, CD11b, IL1β, TNFα and CRP in the brain or plasma (*10*), but did not observe any significant differences that were associated with the activity levels in the OFA (Supplementary Table 6). Brain cholesterol levels, which were measured by lipidomic analysis (*10*), were lower in apoE2-TR mice, and higher in *Apoe*-KO mice, compared to apoE3-TR or apoE4-TR mice (Fig. 4I). Interestingly, brain cholesterol levels were inversely correlated with apoE levels in the brain (Fig. 4J) and the activity levels in the OFA (Fig. 4K). Among plasma cholesterols, only HDL-cholesterol levels, which were lowest in *Apoe*-KO mice (Fig. 4L) were associated with the activity levels in the OFA across *APOE* genotypes (Fig. 4M and Supplementary Table 6). Notably, plasma triglyceride levels, which were highest in apoE2-TR mice (Fig. 4N), were also associated with the activity levels in the OFA (Fig. 4O and Supplementary Table 6) and plasma apoE levels (Fig. 4P). These plasma lipid profiles showed trends toward age-associated decreases in HDL-cholesterol levels in *Apoe*-KO mice, and triglyceride levels in apoE4-TR and apoE3-TR mice. Notably, those age-associated decreases were not observed in apoE2-TR mice (Fig. 4Q&R). These animal data suggest that distinct apoE isoform levels, where *APOE2* has the highest, and associated lipid metabolism, might be involved in the beneficial effects of *APOE2* on longevity phenotypes.

**Fig. 4.**
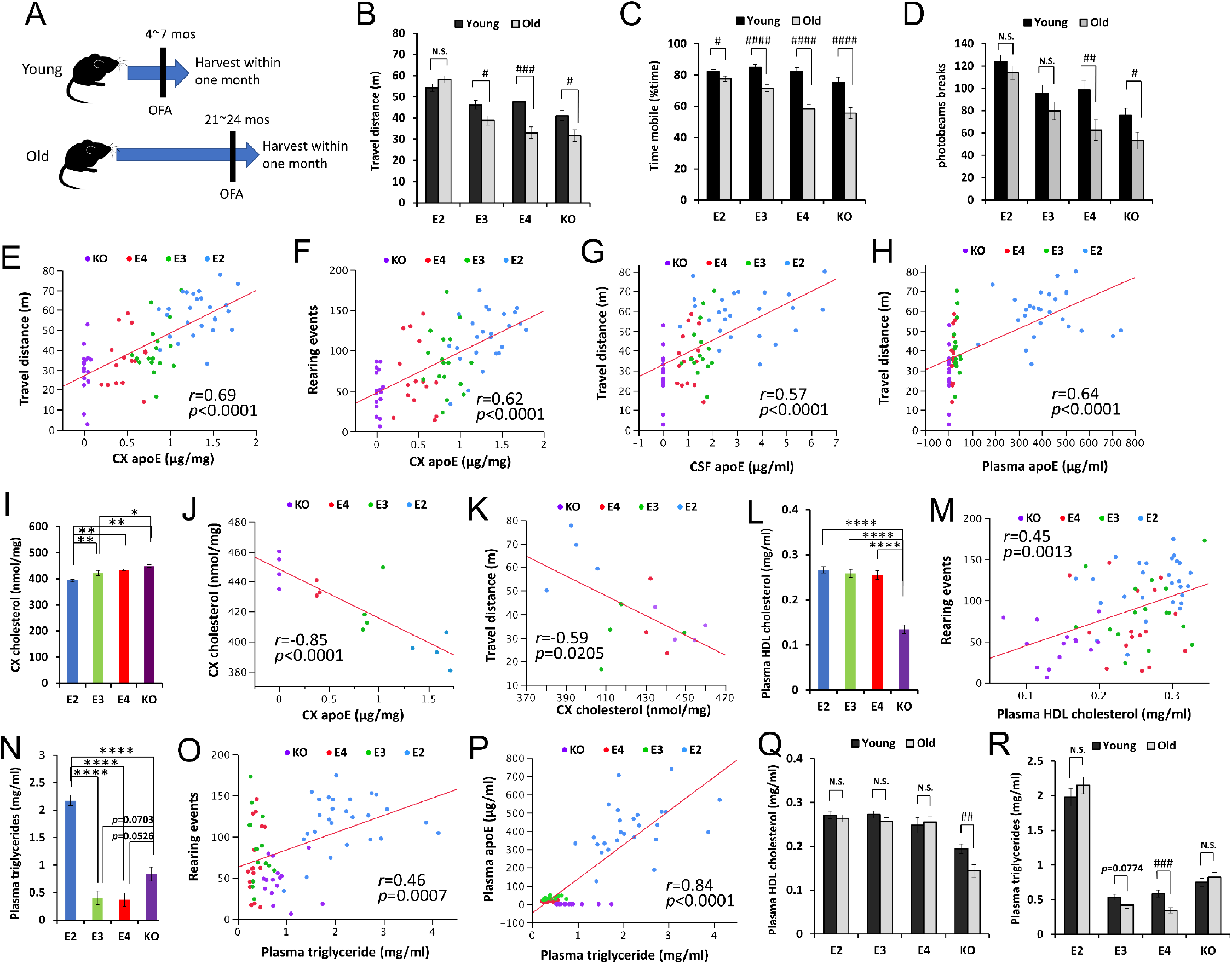
ApoE and lipid levels correlate with activity. (A) The timeline of biochemically assessed cohort. The OFA was performed within one month before harvest. (B-D) Total distance traveled (B), percent time spent mobile (C), and number of rearing events (D) were compared between young and old mice within the same *APOE* genotype after adjusting for sex (n = 16-28 mice/group). (E-H) ApoE levels in the brain cortex (CX, E & F), CSF (G) and plasma (H) are plotted against the total distance traveled (E, G & H), or number of rearing events in the OFA (F). (I) Levels of cholesterol in the brain cortex of old male mice were compared among *APOE* genotypes (n = 3-4 mice/group) and are plotted against apoE levels in the cortex (J) and total distance traveled in the OFA (K). (L-R) Plasma HDL-cholesterol levels (L & M) or triglyceride levels (N-P) were compared among *APOE* genotypes in old mice (L & N) or between young and old mice within the same *APOE* genotype (Q & R) after adjusting for sex and plotted against the number of rearing events in the OFA (M & O) or plasma apoE levels (P) (n = 14-27 mice/group). Data are presented as adjusted means ± standard errors of the means. *p<0.05, **p<0.01, ***p<0.001, ****p<0.0001; compared among *APOE* genotypes using the Tukey-Kramer test (I), Tukey’s HSD test (L & N), or #p<0.05, ##p<0.01, ###p<0.001, ####p<0.0001; compared between young and old mice using two-sided Student’s t test (B-D, Q & R). (E-H, J, K, M, O & P) Correlation coefficients (r) and p-values were calculated using the Pearson correlation test. N.S. = not significant.

## Discussion

Despite strong evidence that *APOE2* contributes to longevity, the underlying mechanism is not clear. Because *APOE2* and *APOE4* respectively reduces and increases the risk of AD, there is a critical need to address whether the effects of *APOE* genotype on longevity are mediated through such AD risks. In this study, we uniquely combined human and mouse model studies and observed that *APOE2* can contribute to longevity independent of AD risks.

Although many clinical studies have observed that *APOE* is associated with longevity, no studies have succeeded in fully distinguishing the longevity effects of *APOE* from its AD effects. NACC database contains both clinical and neuropathological data of large number of AD patients as well as cognitively normal subjects with *APOE* genotypes. Such peculiarities of NACC database indeed allowed us to evaluate whether *APOE-*associated longevity is related to AD or not. While we observed significant *APOE* effects on mortality by adjusting cognitive status, or by stratifying AD and non-AD status, we did not observe significant *APOE* effects in cognitively normal individuals, although there are some trends of significance (data not shown). One possible reason is that compared to demented subjects, there is a smaller number of death event in cognitively normal individuals in NACC database, making it difficult to assess their survival. Another possible reason is that as *APOE2* can also protect against age-related cognitive decline independent of AD (*10*), cognitive decline could be a downstream event of *APOE*-associated pathways related to longevity independent of AD. Longer follow-up would be necessary to address such possibilities.

To strengthen our conclusion regarding NACC clinical data analyses, we analyzed pure animal models to address human *APOE2* effects on longevity. While we observed significant beneficial effects of *APOE2* on lifespan, compared to *APOE3, APOE4* and *Apoe*-KO, we did not observe a significant difference between *APOE4* versus *APOE3* in the overall survival period (Fig. 2B). Although we observed some harmful effects of *APOE4* on lifespan, as observed in “the early survival” and median age at death, which many previous longevity studies primarily evaluate (*16*), further studies would be needed to confirm that *APOE4* has indeed worse impacts on overall lifespan, compared to *APOE3*, as observed in clinical data. Nonetheless, this is the first study showing the effect of *APOE*, especially for the beneficial effects of *APOE2*, on longevity in animal models in the absence of remarkable amyloid or tau pathology like AD.

While we obtained evidence that preserved activity is associated with *APOE2*-related longer lifespan from both clinical and animal studies, similar phenomena were indeed observed in other models of longevity. Calorie restriction, the most validated method to promote longevity in various organisms, was observed to enhance or preserve physical activity, spontaneous locomotive activity or voluntary wheel running activity (*25-28*). Notably, Ingram et al observed in aged C57BL/6J mice, while exploratory activity was associated with lifespan, similar to our findings, other forms of physical activity, such as the scores of rotarod tests, were not (*29*), consistent with this study. Although it is not clear whether enhanced spontaneous or voluntary activity is directly involved in prolonged lifespan (*30*), it would be a potential surrogate maker reflecting longer lifespan.

In this line, significant associations between the levels of activity and those of apoE protein and related lipids (Fig. 4), provide important insights into *APOE* effects on longevity. While OFA activities associate with apoE levels in both CNS and periphery, its associations with cholesterols depends on its subtypes: OFA activities associated with total cholesterols in the brain, but not plasma, whereas they associated with HDL-cholesterol levels in the plasma. These effects maybe mediated by different metabolism of apoE and cholesterols in the brain and periphery as well as distinct effects of absence/presence of each apoE isoforms (*1*). Previous studies have shown that higher HDL-cholesterol levels in plasma were associated with longer lifespan, although effects of plasma total cholesterols were contradictory (*31-34*), likely supporting our results. While *APOE2* carriers have increased levels of plasma triglycerides (*35*), the relationship among apoE2, triglycerides and longevity has not been addressed. Recent studies demonstrate that triglycerides can promote longevity in flies and yeasts (*28, 36*). Moreover, such studies also observed that triglyceride levels are associated with more activity (*28*). Thus, higher HDL-cholesterol and triglyceride levels regulated by apoE2 might both contribute to *APOE2*-mediated longevity although further study is necessary.

There are potential limitations of our study; 1) NACC is not a population-based study, but a referral-or volunteer-based study, making it inappropriate to estimate the life expectancy of general population, 2) in NACC database, despite a longitudinal design, many participants were followed in a short period, 3) cause of death are unavailable, and 4) we did not consider all potential confounding factors associated with death. Thus, we also have conducted additional study using pure animal models expressing each human apoE isoforms, which can address *APOE* effects on longevity in the absence of such potential limitations associated with clinical data. Though additional clinical studies would be warranted, it would be difficult for an individual cohort study when considering lower frequency of *APOE2* carrier as well as general difficulty to combine survival analysis and neuropathological assessment. Also, objective evaluation of activity status would be necessary to confirm our findings regarding preserved activity of *APOE2* carrier in GDS test.

In summary, this study provides complementary evidence from clinical and preclinical data that *APOE2* promotes longevity independent of AD. Our results also indicate that preserved activity during aging might be a good surrogate marker underlying *APOE2*-associated longevity. Moreover, our animal experiments suggest that apoE protein levels, where *APOE2* shows the highest, and associated lipid metabolism might play important roles in *APOE2*-associated longevity. As *APOE2* is a leading longevity gene, understanding the related mechanisms should provide critical insights into healthy living and longevity in our aging society.

## General

We thank Ms. Nancy N. Diehl, Ms. Sarah E. Monsell, Ms. Lilah M. Besser, Ms. Merilee A. Teylan, and Mr. Zachary Miller for assisting with the analysis of the NACC clinical records, Mr. Michael S. Penuliar for assisting with the behavioral experiments, Drs. Mary Jo Ladu and Leon M. Tai for providing the apoE ELISA protocol, Dr. Miao Wang for assisting with the lipidomics study, Dr. Douglas C. Page, Ms. Theresa G. Stile and other Mayo animal facility staff members for caring for the animals, Olivia N. Attrebi for carefully reading this manuscript, and Dr. Yu Yamazaki and Dr. Naoyuki Sato for kindly providing assistance and participating in discussions. The NACC contributors are described in Supplementary Table 7.

## Author contributions

Mi.S., T.K., J.D.F., M.G.H., and G.B. contributed to the concept and study design. Mi.S., T.K., M.T., N.N.D., A.K., Mo.S., Y.F., J.Z., X.H., P.M.S., G.W.R., J.D.F., M.G.H., and G.B. contributed to data acquisition and analysis. Mi.S., and M.G.H. contributed to drafting the manuscript and figures. All authors edited and reviewed the final manuscript.

## Data and materials availability

Data from these experiments are available from the corresponding author upon reasonable request. The clinical data are available from <u>NACC</u> upon request.

