## Supplementary for "*APOE2* promotes longevity independent of Alzheimer’s disease"

**Supplementary Table 1: Subject characteristics in the overall NACC cohort**

| Variable | All subjects (N=24,661) | *APOE3* (ε3/ε3) subjects (N=12,387) | *APOE2* (ε2/ε2 or ε2/ε3) subjects (N=2,269) | *APOE4* (ε3/ε4 or ε4/ε4) subjects (N=9,354) | ε2/ε4 subjects (excluded, N=651) |
| --- | --- | --- | --- | --- | --- |
| Age at initial visit (years) | 73 (18, 109) | 73 (18, 109) | 73 (20, 102) | 72 (20, 99) | 73 (22, 100) |
| Age at final visit (years) | 76 (18, 110) | 76 (18, 110) | 77 (20, 105) | 75 (20, 106) | 76 (22, 103) |
| Sex (female) | 13,882 (56.3%) | 7,016 (56.6%) | 1,294 (57.0%) | 5,185 (55.4%) | 387 (59.4%) |
| Race |  |  |  |  |  |
| White | 20,753 (84.2%) | 10,552 (85.2%) | 1,845 (81.3%) | 7,854 (84.0%) | 502 (77.1%) |
| Black | 2,887 (11.7%) | 1,221 (9.9%) | 343 (15.1%) | 1,187 (12.7%) | 136 (20.9%) |
| Other | 1,021 (4.1%) | 614 (5.0%) | 81 (3.6%) | 313 (3.3%) | 13 (2.0%) |
| Cognitive status at final visit |  |  |  |  |  |
| Normal cognition | 8,869 (36.0%) | 5,187 (41.9%) | 1,085 (47.8%) | 2,369 (25.3%) | 228 (35.0%) |
| Impaired but not MCI | 1,026 (4.2%) | 598 (4.8%) | 126 (5.6%) | 277 (3.0%) | 25 (3.8%) |
| MCI | 3,853 (15.6%) | 2,009 (16.2%) | 384 (16.9%) | 1,349 (14.4%) | 111 (17.1%) |
| Dementia | 10,913 (44.3%) | 4,593 (37.1%) | 674 (29.7%) | 5,359 (57.3%) | 287 (44.1%) |
| Clinically diagnosed AD at final visit | 8,704 (35.3%) | 3,325 (26.8%) | 449 (19.8%) | 4,685 (50.1%) | 245 (37.6%) |
| CV risk factors (present at any visit) | 76(18, 110) | 76 (18, 110) | 77 (20, 105) | 75 (20, 106) | 76 (22, 103) |
| Hypertension | 14,164 (57.5%) | 7,182 (58.0%) | 1,368 (60.4%) | 5,234 (56.0%) | 380 (58.4%) |
| Transient ischemic attack | 1,729 (7.0%) | 898 (7.3%) | 174 (7.7%) | 610 (6.6%) | 47 (7.3%) |
| Pacemaker | 1,103 (4.5%) | 552 (4.5%) | 121 (5.4%) | 406 (4.4%) | 24 (3.8%) |
| Angioplasty/ endarterectomy/stent | 2,071 (8.4%) | 1,064 (8.6%) | 187 (8.2%) | 764 (8.2%) | 56 (8.6%) |
| Heart attack/cardiac arrest | 1,798 (7.3%) | 939 (7.6%) | 173 (7.6%) | 651 (7.0%) | 35 (5.4%) |
| Cardiac bypass procedure | 1,260 (5.1%) | 658 (5.3%) | 104 (4.6%) | 472 (5.0%) | 26 (4.0%) |
| Atrial fibrillation | 2,494 (10.1%) | 1,331 (10.8%) | 247 (10.9%) | 851 (9.1%) | 65 (10.0%) |
| Hypercholesterolemia | 14,443 (58.8%) | 7,215 (58.5%) | 1,000 (44.3%) | 5,919 (63.5%) | 309 (47.5%) |
| Congestive heart failure | 1,021 (4.1%) | 569 (4.6%) | 127 (5.6%) | 302 (3.2%) | 23 (3.5%) |
| Stroke | 1,707 (6.9%) | 851 (6.9%) | 218 (9.6%) | 592 (6.3%) | 46 (7.1%) |
| Death | 5,413 (21.9%) | 2,624 (21.2%) | 444 (19.6%) | 2,196 (23.5%) | 149 (22.9%) |
| With Neuropathological assessment (%Death) | 3,528 (65.2%) | 1,700 (64.8%) | 282 (63.5%) | 1,452 (66.1%) | 94 (63.1%) |
| The sample median (minimum, maximum) is given for continuous variables. Information was unavailable regarding hypertension (N=28), transient ischemic attack (N=116), pacemaker (N=260), angioplasty/endarterectomy/stent (N=19), heart attack/cardiac arrest (N=42), cardiac bypass procedure (N=9), atrial fibrillation (N=48), hypercholesterolemia (N=102), congestive heart failure (N=25), and stroke (N=45). | | | | | |

**Supplementary Table 2: Subject characteristics for individuals in the NACC cohort with a neuropathological assessment**

| Variable | All subjects (N=3,528) | *APOE3* (ε3/ε3) subjects (N=1,700) | *APOE2* (ε2/ε2 or ε2/ε3) subjects (N=282) | *APOE4* (ε3/ε4 or ε4/ε4) subjects (N=1,452) | ε2/ε4 subjects (N=94) |
| --- | --- | --- | --- | --- | --- |
| Age at initial visit (years) | 78 (26, 109) | 80 (26, 109) | 80 (38, 102) | 76 (31, 99) | 80 (49, 100) |
| Age at final visit (years) | 81 (26, 110) | 83 (26, 110) | 83 (38, 105) | 79 (31, 105) | 83 (52, 101) |
| Sex (female) | 1617 (45.8%) | 795 (46.8%) | 136 (48.2%) | 636 (43.8%) | 50 (53.2%) |
| Race |  |  |  |  |  |
| White | 3362 (95.3%) | 1634 (96.1%) | 269 (95.4%) | 1368 (94.2%) | 91 (96.8%) |
| Black | 120 (3.4%) | 38 (2.2%) | 11 (3.9%) | 69 (4.8%) | 2 (2.1%) |
| Other | 46 (1.3%) | 28 (1.6%) | 2 (0.7%) | 15 (1.0%) | 1 (1.1%) |
| Cognitive status at final visit |  |  |  |  |  |
| Normal cognition | 406 (11.5%) | 269 (15.8%) | 64 (22.7%) | 66 (4.5%) | 7 (7.4%) |
| Impaired but not MCI | 52 (1.5%) | 30 (1.8%) | 6 (2.1%) | 15 (1.0%) | 1 (1.1%) |
| MCI | 306 (8.7%) | 183 (10.8%) | 42 (14.9%) | 71 (4.9%) | 10 (10.6%) |
| Dementia | 2764 (78.3%) | 1218 (71.6%) | 170 (60.3%) | 1300 (89.5%) | 76 (80.9%) |
| Clinically diagnosed AD at final visit | 2046 (58.0%) | 824 (48.5%) | 101 (35.8%) | 1057 (72.8%) | 64 (68.1%) |
| CV risk factors (present at any visit) |  |  |  |  |  |
| Hypertension | 2044 (58.0%) | 1027 (60.4%) | 179 (63.5%) | 787 (54.3%) | 51 (54.3%) |
| Transient ischemic attack | 349 (10.0%) | 178 (10.5%) | 32 (11.4%) | 127 (8.8%) | 12 (12.9%) |
| Pacemaker | 260 (7.4%) | 140 (8.2%) | 27 (9.6%) | 89 (6.1%) | 4 (4.3%) |
| Angioplasty/ endarterectomy/stent | 367 (10.4%) | 178 (10.5%) | 28 (9.9%) | 150 (10.3%) | 11 (11.7%) |
| Heart attack/cardiac arrest | 393 (11.2%) | 214 (12.6%) | 28 (9.9%) | 144 (9.9%) | 7 (7.4%) |
| Cardiac bypass procedure | 264 (7.5%) | 139 (8.2%) | 20 (7.1%) | 100 (6.9%) | 5 (5.3%) |
| Atrial fibrillation | 549 (15.6%) | 288 (17.0%) | 41 (14.5%) | 205 (14.1%) | 15 (16.0%) |
| Hypercholesterolemia | 1879 (53.5%) | 899 (53.1%) | 115 (40.9%) | 829 (57.5%) | 36 (38.3%) |
| Congestive heart failure | 322 (9.1%) | 189 (11.1%) | 38 (13.5%) | 89 (6.1%) | 6 (6.4%) |
| Stroke | 401 (11.4%) | 203 (12.0%) | 48 (17.0%) | 141 (9.7%) | 9 (9.7%) |
| Density of neocortical neuritic plaques CERAD score |  |  |  |  |  |
| No neuritic plaques | 771 (21.9%) | 520 (30.7%) | 138 (49.1%) | 100 (6.9%) | 13 (13.8%) |
| Sparse neuritic plaques | 480 (13.7%) | 238 (14.0%) | 50 (17.8%) | 177 (12.2%) | 15 (16.0%) |
| Moderate neuritic plaques | 660 (18.8%) | 350 (20.7%) | 39 (13.9%) | 250 (17.3%) | 21 (22.3%) |
| Frequent neuritic plaques | 1605 (45.6%) | 586 (34.6%) | 54 (19.2%) | 920 (63.6%) | 45 (47.9%) |
| Density of diffuse plaques CERAD score |  |  |  |  |  |
| No diffuse plaques | 513 (16.0%) | 346 (22.6%) | 96 (37.5%) | 64 (4.8%) | 7 (8.0%) |
| Sparse diffuse plaques | 457 (14.3%) | 251 (16.4%) | 50 (19.5%) | 144 (10.8%) | 12 (13.8%) |
| Moderate diffuse plaques | 525 (16.4%) | 253 (16.5%) | 40 (15.6%) | 225 (16.9%) | 7 (8.0%) |
| Frequent diffuse plaques | 1705 (53.3%) | 679 (44.4%) | 70 (27.3%) | 895 (67.4%) | 61 (70.1%) |
| Braak NFT stage |  |  |  |  |  |
| 0 | 196 (5.6%) | 134 (8.0%) | 32 (11.5%) | 28 (1.9%) | 2 (2.1%) |
| I | 301 (8.6%) | 178 (10.6%) | 51 (18.3%) | 69 (4.8%) | 3 (3.2%) |
| II | 406 (11.6%) | 266 (15.8%) | 49 (17.6%) | 78 (5.4%) | 13 (13.8%) |
| III | 324 (9.3%) | 172 (10.2%) | 44 (15.8%) | 98 (6.8%) | 10 (10.6%) |
| IV | 494 (14.1%) | 246 (14.6%) | 47 (16.8%) | 183 (12.7%) | 18 (19.1%) |
| V | 639 (18.3%) | 254 (15.1%) | 21 (7.5%) | 341 (23.6%) | 23 (24.5%) |
| VI | 1113 (31.8%) | 413 (24.6%) | 34 (12.2%) | 641 (44.4%) | 25 (26.6%) |
| Vascular pathology | 3384 (97.3%) | 1601 (96.2%) | 265 (96.4%) | 1425 (98.8%) | 93 (98.9%) |
| Minimal amyloid pathology | 592 (16.8%) | 409 (24.1%) | 117 (41.6%) | 58 (4.0%) | 8 (8.5%) |
| The sample median (minimum, maximum) is given for continuous variables. Information was unavailable regarding hypertension (N=3), transient ischemic attack (N=24), pacemaker (N=1), heart attack/cardiac arrest (N=4), atrial fibrillation (N=3), hypercholesterolemia (N=17), congestive heart failure (N=4), stroke (N=7), .density of neocortical neuritic plaques CERAD score (N=12), density of diffuse plaques CERAD score (N=328), Braak NFT stage (N=55), presence of vascular pathology (N=51), and minimal amyloid pathology. | | | | | |

**Supplementary Table 3: Association between *APOE* genotype and survival using the NACC data**

|  | N | Survival at age 90, % (95% CI) | Adjusting for sex and race | | Adjusting for sex, race, cognitive status at final visit, and AD at final visit^1^ | | Adjusting for sex, race, cognitive status at final visit, AD at final visit^2^, and CV factors^3^ | |
| --- | --- | --- | --- | --- | --- | --- | --- | --- |
| ***APOE* genotype** |  |  | HR (95% CI) | P-value | HR (95% CI) | P-value | HR (95% CI) | P-value |
| All subjects (N=24,661) |  |  |  |  |  |  |  |  |
| *APOE3* (ε3/ε3) | 12,387 | 57.2 (55.5, 59.0) | 1.00 (reference) | N/A | 1.00 (reference) | N/A | 1.00 (reference) | N/A |
| *APOE2* (ε2/ε2 or ε2/ε3) | 2,269 | 64.8 (61.2, 68.7) | 0.84 (0.76, 0.93) | 0.0005 | 0.87 (0.78, 0.96) | 0.005 | 0.88 (0.80, 0.98) | 0.014 |
| *APOE4* (ε3/ε4 or ε4/ε4) | 9,354 | 40.2 (38.0, 42.5) | 1.52 (1.44, 1.61) | <0.0001 | 1.36 (1.28, 1.44) | <0.0001 | 1.34 (1.27, 1.43) | <0.0001 |
| ε2/ε4 | 651 | 43.5 (36.0, 52.5) | 1.24 (1.05, 1.46) | 0.010 | 1.15 (0.98, 1.36) | 0.091 | 1.12 (0.95, 1.33) | 0.17 |
| No AD at final visit (15,957) |  |  |  |  |  |  |  |  |
| *APOE3* (ε3/ε3) | 9,062 | 63.3 (61.1, 65.5) | 1.00 (reference) | N/A | N/A | N/A | 1.00 (reference) | N/A |
| *APOE2* (ε2/ε2 or ε2/ε3) | 1,820 | 67.5 (63.1, 72.2) | 0.88 (0.77, 1.00) | 0.052 | N/A | N/A | 0.86 (0.76, 0.98) | 0.028 |
| *APOE4* (ε3/ε4 or ε4/ε4) | 4,669 | 56.2 (52.3, 60.3) | 1.24 (1.13, 1.37) | <0.0001 | N/A | N/A | 1.26 (1.15, 1.39) | <0.0001 |
| ε2/ε4 | 406 | 56.9 (44.8, 72.2) | 1.03 (0.78, 1.36) | 0.83 | N/A | N/A | 1.03 (0.78, 1.36) | 0.84 |
| AD at final visit (N=8,704) |  |  |  |  |  |  |  |  |
| *APOE3* (ε3/ε3) | 3,325 | 49.4 (46.9, 52.0) | 1.00 (reference) | N/A | N/A | N/A | 1.00 (reference) | N/A |
| *APOE2* (ε2/ε2 or ε2/ε3) | 449 | 58.3 (52.2, 65.2) | 0.84 (0.71, 0.99) | 0.033 | N/A | N/A | 0.84 (0.71, 0.99) | 0.036 |
| *APOE4* (ε3/ε4 or ε4/ε4) | 4,685 | 33.2 (30.7, 35.8) | 1.47 (1.36, 1.59) | <0.0001 | N/A | N/A | 1.43 (1.33, 1.53) | <0.0001 |
| ε2/ε4 | 245 | 34.8 (26.4, 46.1) | 1.28 (1.04, 1.57) | 0.022 | N/A | N/A | 1.24 (1.00, 1.53) | 0.050 |
| Neuropathological assessment (N=3528)^4^ |  |  |  |  |  |  |  |  |
| *APOE3* (ε3/ε3) | 1700 | 24.1 (22.2, 26.2) | 1.00 (reference) | N/A | N/A | N/A | 1.00 (reference) | N/A |
| *APOE2* (ε2/ε2 or ε2/ε3) | 282 | 31.9 (2.9, 37.9) | 0.83 (0.72, 0.95) | 0.006 | N/A | N/A | 0.88 (0.76, 1.01) | 0.062 |
| *APOE4* (ε3/ε4 or ε4/ε4) | 1452 | 11.2 (9.7, 12.9) | 1.46 (1.34, 1.58) | <0.0001 | N/A | N/A | 1.41 (1.30, 1.53) | <0.0001 |
| ε2/ε4 | 94 | 20.2 (13.5, 20.2) | 1.06 (0.85, 1.32) | 0.60 | N/A | N/A | 0.90 (0.72, 1.12) | 0.33 |
| Minimal amyloid pathology (N=592) |  |  |  |  |  |  |  |  |
| *APOE3* (ε3/ε3) | 409 | 19.6 (16.1, 23.8) | 1.00 (reference) | N/A | N/A | N/A | 1.00 (reference) | N/A |
| *APOE2* (ε2/ε2 or ε2/ε3) | 117 | 28.2 (21.1, 37.7) | 0.83 (0.73, 0.94) | 0.003 | N/A | N/A | 0.78 (0.63, 0.96) | 0.021 |
| *APOE4* (ε3/ε4 or ε4/ε4) | 58 | 13.8 (7.2, 26.2) | 1.52 (1.41, 1.63) | <0.0001 | N/A | N/A | 1.36 (1.03, 1.82) | 0.033 |
| ε2/ε4 | 8 | 25.0 (7.5, 83.0) | 1.01 (0.82, 1.25) | 0.90 | N/A | N/A | 1.02 (0.50, 2.09) | 0.95 |
| HR=hazard ratio; CI=confidence interval; AD=Alzheimer’s disease; CV=cardiovascular. HRs, 95% CIs, and p-values result from Cox proportional hazards regression models. ^1^ These analyses were only performed when considering all subjects. ^2^ Cognitive status at final visit and AD at final visit were adjusted for only in the model involving all subjects. ^3^ CV factors included hypertension, transient ischemic attack, pacemaker, angioplasty/endarterectomy/stent, heart attack/cardiac arrest, atrial fibrillation, hypercholesterolemia, congestive heart failure, and stroke. ^4^ For the neuropathological assessment cohort, all models were additionally adjusted for CERAD diffuse plaque score, CERAD neuritic plaque score, Braak NFT stage, and the presence of vascular pathology. P-values < 0.025 are considered as statistically significant after applying a Bonferroni correction for multiple testing for the two primary comparisons of survival that were made (i.e. between *APOE3* and *APOE4* subjects, and between *APOE3* and *APOE2* subjects). | | | | | | | | |

**Supplementary Table 4: Summary of mouse cohorts**

|  | **ApoE2-TR** | **ApoE3-TR** | **ApoE4-TR** | ***Apoe*-KO** | ***p*-value** |
| --- | --- | --- | --- | --- | --- |
| **Survival cohort** | | | | | |
| **Total No.** | 29 | 27 | 34 | 28 | — |
| **Female No. (%)** | 17 (58.6) | 14 (51.9) | 20 (58.8) | 14 (50.0) | 0.8625 |
| **Age at behavior-tested at old age, month** | 23.5 ± 1.4 [21.1-24.6] | 23.8 ± 0.3 [23.3-24.1] | 24.1 ± 0.1 [23.9-24.3] | 22.5 ± 0.9 [21.8-23.9] | <0.0001 |
| **Median age at death, day [95%lower, higher]** | 911 [780-949] | 825 [783-853] | 752.5 [696-796] | 737.5 [660-817] | 0.0041 |
| **Found dead No. (%)** | 14 (48.3) | 19 (70.3) | 24 (70.6) | 21 (75.0) | 0.1413 |
| **Euthanized No. (%)** | 15 (51.7) | 8 (29.6) | 10 (29.4) | 7 (25.0) |  |
| **Hunched, weight loss, lethergic (% of euthanized)** | 3 | 6 | 5 | 4 | 0.0653 |
| **Tumor (% of euthanized)** | 2 | 2 | 3 | 0 | 0.2549 |
| **Recurrent UD, skin problem (% of euthanized)** | 5 | 0 | 2 | 2 | 0.3086 |
| **Prolapse (% of euthanized)** | 1 | 0 | 0 | 0 | 0.6448 |
| **Multiple (% of euthanized)** | 2 | 0 | 0 | 1 | 0.4415 |
| **Other reasons (due to end of study)** | 2 | 0 | 0 | 0 | 0.3323 |
| **Biochemically assessed cohort** | | | | | |
| **Young cohort No.** | 28 | 26 | 26 | 23 | — |
| **Female No. (%)** | 14 (50.0) | 12 (46.2) | 14 (53.8) | 11 (47.8) | 0.9520 |
| **Age at sacrifice, month** | 7.3 ± 0.7 [6.3-7.9] | 7.6 ± 0.2 [7.4-7.8] | 7.2 ± 0.3 [6.6-7.5] | 6.2 ± 0.8 [5.2-7.4] | <0.0001 |
| **Male body weight, gram** | 28.7 ± 2.1 [25.9-33.1] | 29.5 ± 2.7 [22.2-33.4] | 30.5 ± 2.8 [25.3-34.9] | 29.2 ± 2.4 [23.4-32.6] | 0.3432 |
| **Female body weight, gram** | 24.7 ± 2.5 [21.7-28.9] | 21.9 ± 1.9 [19.4-24.7] | 23.5 ± 1.7 [21.2-27.1] | 24.0 ± 2.6 [19.0-28.2] | 0.0190 |
| **Old cohort No.** | 27 | 17 | 16 | 17 | — |
| **Female No. (%)** | 10 (37.0) | 8 (47.1) | 10 (62.5) | 6 (35.3) | 0.3424 |
| **Age at sacrifice, month** | 22.8 ± 1.0 [21.3-24.7] | 22.6 ± 1.0 [21.7-24.5] | 22.4 ± 0.2 [22.2-22.7] | 22.8 ± 0.6 [21.7-24.1] | 0.3565 |
| **Male body weight, gram** | 31.6 ± 2.3 [27-36.9] | 33.5 ± 3.4 [28.8-38.4] | 32.7 ± 2.1 [30.4-35.8] | 29.5 ± 3.4 [20.2-32.8] | 0.0263 |
| **Female body weight, gram** | 25.1 ± 3.4 [19.3-30.0] | 25.2 ± 3.5 [22.6-33.7] | 25.4 ± 3.7 [17.8-29.3] | 26.2 ± 3.2 [22-30.3] | 0.9425 |
| For continuous data, values are means ± SD [range]. P-values were calculated using one-way ANOVA (continuous data), the Pearson’s chi-square test (categorical value) or log-rank test (median age at death). | | | | | |

**Supplementary Table 5: Effects of *APOE* on “Dropped activities and interests” and other GDS items in cognitively normal subjects over 60 years old**

|  |  |  |  | *APOE2* vs. *APOE3* (reference) | | *APOE4* vs. *APOE3* (reference) | |
| --- | --- | --- | --- | --- | --- | --- | --- |
| **GDS item** | *APOE2* (ε2/ε2 or ε2/ε3, N=1,197) | *APOE3* (ε3/ε3, N=5,717) | *APOE4* (ε3/ε4 or ε4/ε4, N=2,616) | OR (95% CI) | P-value | OR (95% CI) | P-value |
| **Primary item** |  |  |  |  |  |  |  |
| Dropped many of your activities and interests | 149 (12.4%) | 830 (14.5%) | 349 (13.4%) | 0.80 (0.66, 0.96) | 0.019 | 0.98 (0.86, 1.13) | 0.82 |
| **Secondary items** |  |  |  |  |  |  |  |
| Satisfied with your life | 1102 (92.1%) | 5190 (90.9%) | 2379 (91.0%) | 1.19 (0.95, 1.49) | 0.14 | 1.00 (0.85, 1.18) | 0.96 |
| Feel that your life is empty | 57 (4.8%) | 310 (5.4%) | 139 (5.3%) | 0.85 (0.64, 1.14) | 0.29 | 1.03 (0.83, 1.27) | 0.80 |
| Often get bored | 93 (7.8%) | 497 (8.7%) | 196 (7.5%) | 0.87 (0.69, 1.09) | 0.22 | 0.84 (0.71, 1.00) | 0.053 |
| In good spirits most of the time | 1134 (94.7%) | 5410 (94.7%) | 2479 (94.8%) | 0.99 (0.75, 1.30) | 0.91 | 1.02 (0.83, 1.26) | 0.85 |
| Afraid something bad happen is going to happen to you | 80 (6.7%) | 437 (7.6%) | 211 (8.1%) | 0.90 (0.70, 1.15) | 0.38 | 1.08 (0.91, 1.28) | 0.39 |
| Feel happy most of the time | 1099 (91.9%) | 5206 (91.3%) | 2388 (91.3%) | 1.07 (0.86, 1.35) | 0.54 | 0.99 (0.84, 1.16) | 0.87 |
| Often feel helpless | 72 (6.0%) | 471 (8.2%) | 180 (6.9%) | 0.69 (0.53, 0.89) | 0.005 | 0.90 (0.75, 1.07) | 0.24 |
| Prefer to stay at home rather than going out and doing new things | 274 (22.9%) | 1376 (24.1%) | 583 (22.3%) | 0.94 (0.81, 1.09) | 0.39 | 0.94 (0.84, 1.05) | 0.24 |
| Feel that you have more problems with memory than most | 126 (10.5%) | 710 (12.4%) | 378 (14.5%) | 0.86 (0.70, 1.05) | 0.13 | 1.18 (1.03, 1.35) | 0.020 |
| Think it is wonderful to be alive now | 1134 (94.7%) | 5433 (95.1%) | 2494 (95.4%) | 0.93 (0.70, 1.24) | 0.62 | 0.93 (0.74, 1.16) | 0.52 |
| Feel pretty worthless the way you are now | 52 (4.3%) | 272 (4.8%) | 91 (3.5%) | 0.88 (0.65, 1.19) | 0.40 | 0.82 (0.64, 1.05) | 0.12 |
| Feel full of energy | 755 (63.1%) | 3717 (65.0%) | 1786 (68.3%) | 0.95 (0.83, 1.08) | 0.40 | 1.07 (0.97, 1.18) | 0.19 |
| Feel that your situation is hopeless | 23 (1.9%) | 174 (3.0%) | 69 (2.6%) | 0.62 (0.40, 0.96) | 0.034 | 0.96 (0.72, 1.28) | 0.79 |
| Think that most people are better off than you are | 37 (3.1%) | 175 (3.1%) | 78 (3.0%) | 1.00 (0.70, 1.44) | 1.00 | 0.99 (0.75, 1.30) | 0.92 |
| Total GDS score | 1.44 (0, 2) | 1.55 (0, 2) | 1.46 (0, 2) | 0.92 (0.82, 1.04) | 0.17 | 0.99 (0.91, 1.07) | 0.73 |
| ORs, 95% CIs, and p-values result from binary logistic regression models for analysis of individual GDS items, and from proportional odds logistic regression models for analysis of the ordinal total GDS score, which is described as the mean (25^th^ percentile, 75^th^ percentile). Information was unavailable for GDS items for a maximum of 23 subjects. | | | | | | | |

**Supplementary Table 6: Associations between activity levels in the OFA and protein/lipids levels across *APOE* genotype groups at old age**

|  | **Travel distance** | | **Time mobile** | | **Rearing events** | |
| --- | --- | --- | --- | --- | --- | --- |
|  | ***r*** | ***p*-value** | ***r*** | ***p*-value** | ***r*** | ***p*-value** |
| **CX apoE** | 0.69 | <0.0001 | 0.59 | <0.0001 | 0.62 | <0.0001 |
| **HP apoE** | 0.68 | <0.0001 | 0.57 | <0.0001 | 0.56 | <0.0001 |
| **CSF apoE** | 0.57 | <0.0001 | 0.42 | 0.0043 | 0.47 | 0.006 |
| **Plasma apoE** | 0.64 | <0.0001 | 0.43 | 0.0018 | 0.54 | <0.0001 |
| **CX PSD95** | -0.15 | 1 | -0.19 | 1 | -0.29 | 0.1630 |
| **HP PSD95** | 0.27 | 0.2658 | 0.28 | 0.2519 | 0.15 | 1 |
| **CX GFAP** | 0.21 | 1 | 0.06 | 1 | -0.09 | 1 |
| **HP GFAP** | 0.14 | 1 | 0.10 | 1 | -0.10 | 1 |
| **CX CD11b** | 0.09 | 1 | 0.06 | 1 | -0.06 | 1 |
| **HP CD11b** | 0.15 | 1 | 0.27 | 0.3072 | -0.07 | 1 |
| **CX IL1β** | 0.24 | 0.5742 | 0.09 | 1 | 0.21 | 1 |
| **CX TNFα** | 0.09 | 1 | -0.07 | 1 | 0.06 | 1 |
| **Plasma CRP** | 0.09 | 1 | 0.137 | 1 | 0.07 | 1 |
| **Plasma total cholesterol** | -0.04 | 1 | -0.24 | 0.6085 | -0.04 | 1 |
| **Plasma HDL cholesterol** | 0.30 | 0.1747 | 0.31 | 0.1338 | 0.45 | 0.0013 |
| **Plasma non-HDL cholesterol** | -0.05 | 1 | -0.25 | 0.5500 | -0.04 | 1 |
| **Plasma triglyceride** | 0.50 | 0.0001 | 0.26 | 0.4152 | 0.46 | 0.0007 |
| Correlation coefficients (r) and p-values were calculated using the Pearson correlation test. P-values were corrected by Bonferroni test adjusted for the number of protein/lipids analyzed in the entire cohort. CX = cortical, HP = hippocampal. | | | | | | |

**Supplementary Table 7:** List of NACC contributors.

| Contributors (PI) | NIA/NIH Funding Resource |
| --- | --- |
| Eric Reiman, MD | P30 AG019610 |
| Neil Kowall, MD | P30 AG013846 |
| Scott Small, MD | P50 AG008702 |
| Allan Levey, MD, PhD | P50 AG025688 |
| Andrew Saykin, PsyD | P30 AG010133 |
| Marilyn Albert, PhD | P50 AG005146 |
| Bradley Hyman, MD, PhD | P50 AG005134 |
| Ronald Petersen, MD, PhD | P50 AG016574 |
| Mary Sano, PhD | P50 AG005138 |
| Steven Ferris, PhD | P30 AG008051 |
| M. Marsel Mesulam, MD | P30 AG013854 |
| Jeffrey Kaye, MD | P30 AG008017 |
| David Bennett, MD | P30 AG010161 |
| Charles DeCarli, MD | P30 AG010129 |
| Frank LaFerla, PhD | P50 AG016573 |
| David Teplow, PhD | P50 AG016570 |
| Douglas Galasko, MD | P50 AG005131 |
| Bruce Miller, MD | P50 AG023501 |
| Russell Swerdlow, MD | P30 AG035982 |
| Linda Van Eldik, PhD | P30 AG028383 |
| John Trojanowski, MD, PhD | P30 AG010124 |
| Oscar Lopez, MD | P50 AG005133 |
| Helena Chui, MD | P50 AG005142 |
| Roger Rosenberg, MD | P30 AG012300 |
| Thomas Montine, MD, PhD | P50 AG005136 |
| Sanjay Asthana, MD, FRCP | P50 AG033514 |
| John Morris, MD | P50 AG005681 |
| Stephen Strittmatter, MD, PhD | P50 AG047270 |
